# Elevated ENO2 Disrupts VSMCs Homeostasis Facilitating the Development of Aortic Dissection

**DOI:** 10.1101/2023.03.23.534041

**Authors:** Weixing Cai, Li Zhang, Jing-Jing Lin, Qiuying Zou, Chaoyun Wang, Xi Yang, Xiaohui Wu, Jianqiang Ye, Hui Zheng, Lin Zhang, Lin Zhong, Xinyao Li, Keyuan Chen, Jiangbin Wu, Liangwan Chen, Yumei Li

**Affiliations:** Department of Cardiovascular Surgery, Fujian Medical University Union Hospital, Fuzhou, China; Fujian Center for Safety Evaluation of New Drug, The School of Pharmacy, Fujian Medical University, Fuzhou, China; Fujian Medical University, Fuzhou, China; Key Laboratory of Cardio-Thoracic Surgery (Fujian Medical University), Fujian Province University, Fuzhou, China; Department of Physiology & Pathophysiology, The School of Basic Medical Sciences, Fujian Medical University, Fuzhou, China; PET/CT center, Fujian Medical University Union Hospital, Fuzhou, China; Department of Cardiac Surgery, The First Affiliated Hospital of Guangzhou Medical University, Guangzhou, China; The School of Pharmacy, Fujian Health College, Fuzhou, China

**Keywords:** Aortic dissection, Autophagy, Enolase 2, Vascular smooth muscle cells

## Abstract

**OBJECTIVE:** Aortic dissection (AD) is a dangerous cardiovascular disease. However, its regulatory mechanism remains poorly understood. Autophagy is an important pathway for maintaining cellular homeostasis. Recent studies have shown that glycolysis and autophagy are essential for the development of AD. Elevated glycolysis can promote cells autophagy, but the relationship between the two is unclear. Enolase 2 (ENO2) is a glycolytic enzyme. Here, we investigated whether ENO2 can regulate autophagy in vascular smooth muscle cells (VSMCs) involved in the progression of AD.

**APPROACH AND RESULTS:** The levels of ENO2 and autophagy-related proteins in aortic tissues from AD patients were identified by qRT-PCR, Western blot and IHC. We found that autophagosome clearance was impaired in aortic tissues and positively correlated with elevated ENO2 expression. Furthermore, aortic application of smooth muscle-specific adeno-associated virus (AAV) inhibiting ENO2 expression attenuated the development of AD in mice. in vitro experiments, downregulation of ENO2 partially restored PDGF-BB-induced impairment of autophagic flux, as evidenced by reduced expression of Beclin-1, SQSTM1, LC3BII/I and increased levels of LAMP2 in human aortic vascular smooth muscle cells (HAVSMCs). Similarly, increased levels of ENO2 exacerbated the autophagic dysfunction in HAVSMCs. Mechanistically, ENO2 may block autophagic flux through the ENO2-GAP43-ATF4 pathway and disrupt cellular homeostasis.

**CONCLUSIONS:** Our data reveal a perturbing role of ENO2 in the homeostasis of VSMCs, suggesting ENO2 as a potential target for intervention in AD.

## Introduction

Aortic dissection (AD) is a fatal cardiovascular disease that arises from an intimal tear, leading to separation of the aortic wall layers, hemorrhagic infiltration, and the risk of aortic rupture^1^. The current clinical treatment of AD is mainly achieved by surgical intervention, and its mortality rate is very high if not treated in time. Therefore, it is crucial to elucidate the pathogenesis of AD.

Enolase, a key glycolytic enzyme, supports a host of functions, including autophagy and apoptosis^2,3^. Specifically, Enolase 2 (ENO2) is not only one of the three enolase isozymes in mammals, but also a multifunctional protein. It is upregulated in many malignancies and may act as a pro-survival factor^4–6^. In the field of cardiovascular medicine, there are few reports on ENO2. Recent studies suggest that ENO2 might be a pathological marker of hypoxia-related cardiovascular remodeling and that dynamic expression can be used to capture the premorbid state of hypoxia-induced pulmonary vascular remodeling^7^. Meanwhile, increased expression of ENO2 can also promote angiogenesis^8^. However, there are few reports on the role of ENO2 in AD.

Autophagy is a process by which cytoplasmic components, such as protein aggregates, are broken down and recycled through the lysosomal apparatus^9^. It achieves cellular homeostasis by selectively degrading harmful protein aggregates and damaged organelles^10^. Accumulating evidence indicates that impaired autophagy contributes to the development of cardiovascular disease^11,12^. The aorta of AD patients may have deregulated autophagy, which is hallmarked by an accumulation of LC3B and increased SQSTM1 protein levels^13–15^. More importantly, the autophagy protein SQSTM1 is considered to be a novel selective target for the prevention and postoperative treatment of aortic aneurysm and aortic dissection. Therefore, further exploration of the underlying mechanisms of impaired autophagy in smooth muscle cells in AD will lay the foundation for future AD prevention and protection.

To this end, a clinical sample study was conducted to discern the association between ENO2 and AD. In addition, an experimental AD model was established using 3-aminopropionitrile fumarate (BAPN) and angiotensin II (Ang II) to further reveal the role of ENO2 in the development of AD and the potential mechanisms that might be involved, providing value for targeting ENO2 for clinical application in AD.

## Materials and methods

### Human aortic samples

We collected abandoned aorta from AD patients after aortic replacement in Fujian Medical University Union Hospital. The diagnosis of AD is confirmed by clinical history, imaging and physical examination. In addition, Normal thoracic aortic tissue was obtained from the heart graft organ donor. None of these patients had cardiovascular disease, and their vessels served as controls. The study was conducted in accordance with the ethical guidelines of the Declaration of Helsinki and approved and conducted under the guidance of the Ethics Committee of Fujian Medical University Union Hospital (No. 2019-36). All subjects signed a written informed consent form. The general information of patients is shown in Table 1. Dissected arteries were collected during the procedure, and part of the tissue was immediately stored in the laboratory at −80℃ for Western blot, while the rest of the tissue was stored in 4% paraformaldehyde for pathological staining and immunofluorescence.

**Table 1.** General information of the patients.

| Group | AD ID | Gender | Age | BP | LVEF | EF% | Diameter |
| --- | --- | --- | --- | --- | --- | --- | --- |
| Control | 1101066 | male | 65 | 93/63 | 37.2 | 67.7% | 28.2 |
|  | 1102136 | male | 37 | 100/70 | 32.5 | 61.4% | 28.3 |
|  | 1104996 | male | 36 | 96/52 | 31.7 | 60.1% | 22.4 |
|  | 1100526 | female | 61 | 106/63 | 38.9 | 70.2% | 26.0 |
|  | 1095362 | female | 53 | 131/97 | 33.9 | 63.7% | 26.1 |
| AD | 1111375 | male | 50 | 178/64 | 37.1 | 67.1% | 40.4 |
|  | 991020 | male | 58 | 181/106 | 47.3 | 78.7% | 45.2 |
|  | 1115681 | male | 71 | 172/78 | 32.1 | 59.8% | 47.7 |
|  | 1114723 | male | 63 | 148/86 | 28.1 | 54.2% | 36.2 |
|  | 1114252 | female | 60 | 96/67 | 39.0 | 70.1% | 51.7 |

### Construction and administration of adeno-associated virus (AAV) serotype 9 vector

To specifically knock down ENO2 in VSMCs, we constructed an AAV9-sm22α-Mir30-m-ENO2-ZsGreen vector (AAV-ENO2) with a smooth muscle-specific promoter SM22α and with zsgreen green fluorescence (Hanbio, Shang Hai, China). An AAV9 expressing GFP (AAV-GFP) with a smooth muscle-specific promoter SM22α and with zsgreen green fluorescence was simultaneously prepared as a control vector.

AAV9 virus was coated in a gel (P2443-1kg, sigma-Aldrich) carrier in the aorta of mice during thoracotomy one week before they were fed drinking water containing 0. 3% BAPN. We have improved the method of adenovirus interference in the thoracic aorta based on the reported literature^16^. By performing open-heart surgery on mice, a gel mixed with adenovirus was applied to the aorta to achieve a better interference effect and also to save the amount of adenovirus.

### Mouse model of AD

The ATVB Council recommends consideration Sex differences in aortic dissection studies, including TAD^17^. In our experiments, we only studied male mice, which are prone to aortic dissection, because male mice have less sex hormone variation^18^. The male mice were fed with 0.3%BAPN (A0408, TCI) in drinking water for 4 weeks. Beginning on the 36th day, Ang II (4474-91-3, Tocris) was injected subcutaneously at a total dose of 1. 44mg/kg twice a day at 6-hour intervals for 3 days. Based on the reported literature we identified the methods used to induce AD ^19^.

### Immunohistochemistry (IHC)

Aortic tissues from patients and mice were embedded in paraffin wax and then cut into 3-μm-thick sections. Paraffin sections were processed in an oven (65°C, 30 min) and then dewaxed by immersion in different concentration gradients of xylene and ethanol. Paraffin sections were processed according to the instructions of the immunohistochemistry kit (UItraSensitive SP mouse/rabbit, MXB), during which the appropriate primary antibody was selected and incubated overnight at 4°C. Staining was performed using the DAB staining kit (DAB-0031, MXB) according to the instructions. Statistics and analysis were performed by Nikon fluorescence microscopy (Nikon ECLIPSE Ni) and Image-Pro Plus 6. 0 (IPP 6. 0). All antibodies used in this study were listed in Supplementary Table 1.

### Aorta Histology

Aortic samples from mice were collected and fixed in 4% paraformaldehyde at room temperature. Aortic tissue samples fixed with paraformaldehyde were embedded in paraffin. Vascular tissue samples were cut into 3-μm sections using a paraffin microtome and stained with hematoxylin-eosin (H&E), Masson trichrome (G1340, Solarbio Life Sciences), and EVG (R20391–3, Yuanye Bio-technology). Mice aortic tissue was observed under a Nikon fluorescence microscope (Nikon ECLIPSE Ni). Statistics and analysis were performed by Nikon fluorescence microscopy (Nikon ECLIPSE Ni) and Image-Pro Plus 6. 0 (IPP 6. 0).

### Cell culture and treatment

HAVSMCs were purchased from Otwo Biotechnology (Shenzhen) Co., Ltd. Cells were cultured with DMEM containing 10% FBS. HAVSMCs (stages 4-6) were inoculated onto plates. cells were treated with different drugs. PDGF-BB 20 ng/mL (P5579-10 μg, Beyotime), recombinant human ENO2 1 μg/mL (P7040-5 μg, Beyotime).

### Immunofluorescence (IF)

Human aortic tissue was embedded in paraffin and cut into 3 µm thick sections. Paraffin sections were processed in the oven (65◦C) and soaked in xylene and alcohol. Alcohol of different concentration gradients was used for dewaxing. The sections were then soaked in 3% H2O2 for 20 min and blocked with goat serum. Then treated with primary antibodies treatment, followed by incubation with anti-mouse secondary antibody, anti-rabbit secondary antibody and DAPI. Statistics and analysis were performed by Nikon fluorescence microscopy (Nikon ECLIPSE Ni) and Image J. All antibodies used in this study were listed in Supplementary Table 1.

HAVSMCs were fixed on glass slides with 4% paraformaldehyde and then treated with appropriate primary antibodies, followed by incubation with appropriate anti-rabbit secondary antibody, anti-mouse secondary antibody and DAPI. Statistics and analysis were performed by Nikon fluorescence microscopy (Nikon ECLIPSE Ni) and Image J. All antibodies used in this study were listed in Supplementary Table 1.

### Plasmids and siRNAs

pcDNA3. 1-GAP43 was ordered from Integrated Biotech Solutions Co., Ltd (Shanghai, China) and transfected with lipo3000 (L3000001-0. 1 mL, Invitrogen) for overexpression of GAP43. To knock down ENO2 and ATF4, ENO2 and ATF4 were purchased from Integrated Biotech Solutions Co., Ltd (Shanghai, China) and were mixed with lipo8000 (C0533-0. 5 mL, Beyotime) for transfection, respectively. siRNA nucleotide sequences are listed in Supplementary Table 2.

### Quantitative real-time polymerase chain reaction (qRT-PCR)

Total RNA from treated cells was isolated using Trizol reagent. mRNA levels were measured using qRT-PCR kit (Takara) in combination with TB Green real-time PCR. Amplification was performed using Applied Biosystems QuanStudio ®5 (Applied Biosystems). Primer sequences are shown in Supplementary Table 3.

### Western blot

Cell and tissue extracts were prepared according to the instructions, and protein concentrations were determined using the BCA Protein Assay Kit (P0010S, Beyotime) with bovine serum albumin (BSA) as the standard. Protein bands were detected by GE AI680 and quantified by Amersham Imager 680 analysis software. All antibodies used in this study were listed in Supplementary Table 1.

### Transmission electron microscopy (TEM) analysis

Specimens were fixed in 2. 5% glutaraldehyde and Milonig’s phosphate buffer (pH=7. 3) from treated HAVSMCs for transmission electron microscopy laboratory examination. Briefly, specimens were washed with Milonig’s phosphate buffer and incubated in 1% osmium tetroxide for 1 hour. The samples were then dehydrated in a series of graded acetone, followed by sample resin immersion, embedding and curing processes. Ultrathin sections (50-100 nm) of the specimens were made with an ultramicrotome and diamond knife. Specimens were double stained with 3% uranyl acetate and lead nitrate, examined and photographed under a transmission electron microscope (Philips, Amsterdam, Netherlands).

### Transfection and evaluation of fluorescent dots

To analyze autophagosomes in HAVSMCs cells, Ad-GFP-LC3 (C3006-1ml, Beyotime) was used according to the manufacturer’s instructions. Briefly, when cells reached 60% confluence, adenovirus was added to the medium. Cells were incubated at 37°C for 12 hours and then transferred to completely fresh medium. After 24 hours of Ad-GFP-LC3 infection, cells were treated with the indicated chemicals or transfected with the indicated siRNAs. Cells were then fixed with 4% formaldehyde for 30 min and stained with DAPI for 10 min at room temperature. Fluorescent images were captured using a Nikon fluorescence microscope (Nikon ECLIPSE Ni). Based on previous studies we determined the timing of infection and the use of Ad-GFP-LC3 ^20^.

### Statistical analysis

The results were expressed as average ± SEM. Two groups of data were compared by double-tailed Student’s t-test, and multiple groups were compared by one-way analysis of variance in GraphPad Prism 8. 0. It is assumed that *p* < 0. 05 indicates statistical significance.

## Results

### The expression of ENO2 was up-regulated in AD patients

We collected aortic tissue. Normal thoracic aortic tissue was obtained from heart transplant organ donors. None of these patients had cardiovascular disease. Aortic tissues were obtained from AD patients after aortic replacement. We excluded AD patients with rare genetic disorders such as Marfan’s syndrome, Loeys-Dietz syndrome. The collected patient information is shown in Table 1. The expression of ENO2 was detected by qRT-PCR and western blot and was verified by IHC assay. It was found that both the mRNA expression and the protein expression of ENO2 were much higher in the aorta tissues of AD patients than those in the normal tissues (Figure 1A-C). Taken together, these data demonstrate increased ENO2 expression in AD lesions in humans. we performed simulations in HAVSMCs. PDGF-BB is commonly used in experimental models of VSMCs pathology ^21^. Following induction with PDGF-BB, HAVSMCs also showed increased expression of ENO2 in HAVSMCs (Figure 1D), suggesting successful establishment of the AD model.

**Figure 1.**
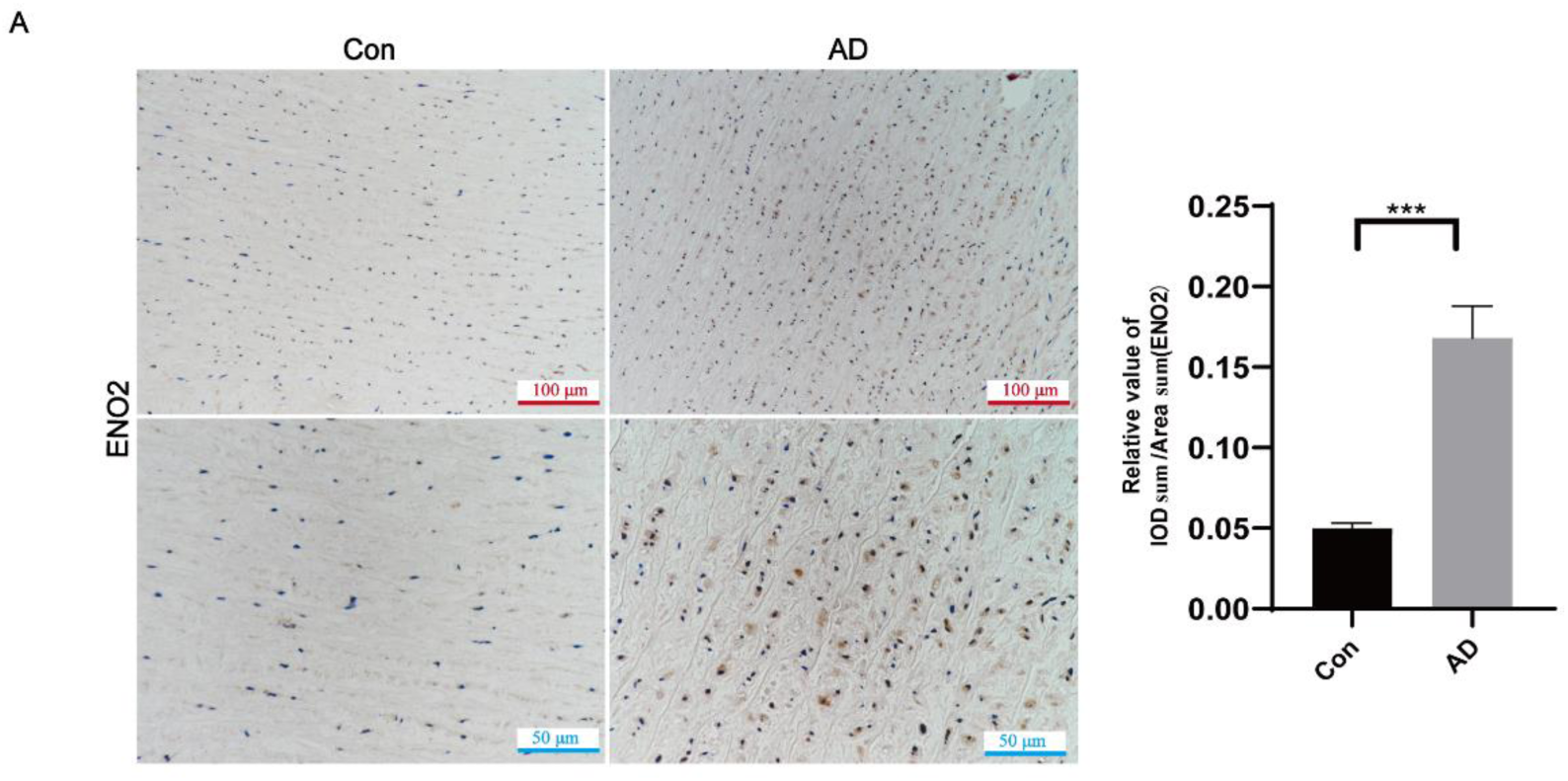
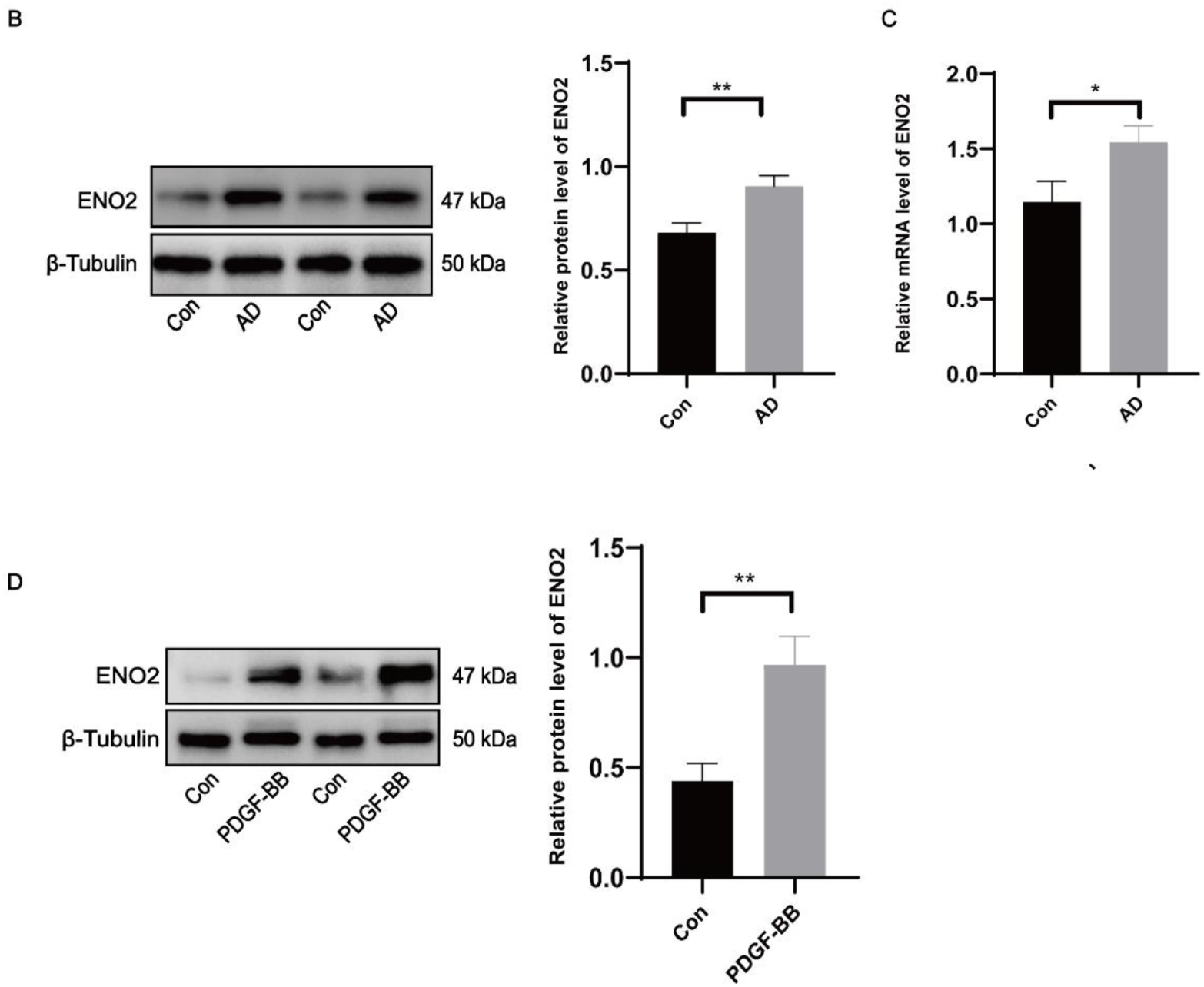
The expression of ENO2 was up-regulated in AD patients. **A** The expression of ENO2 in AD patients and the control group was observed by IHC, n = 5 for each group. **B** The expression of ENO2 in AD patients and the control group was observed by western blot. **C** The mRNA expression of ENO2 in AD patients and the control group was observed by qRT-PCR, n = 5 in each group. **D** ENO2 expression in PDGF-BB-stimulated HAVSMCs, detected by western blot. Data are presented as mean fold-change with error bars for SEM. T-test was used to compare the two groups of data. Statistically significant differences are indicated at * *p* < 0. 05; ** *p* < 0. 01; *** *p* < 0. 001.

### Impaired autophagosome clearance in the aorta of AD patients and a positive correlation with elevated ENO2 expression

To examine the alterations in autophagy in AD patients, we performed western blot to assess the levels of autophagy-related proteins. First, we assayed the levels of LC3B and Beclin-1 in human aorta. LC3B is a marker of autophagy formation, and the ratio of LC3BII to LC3BI is related to the number of autophagosomes^22^. Beclin-1 is also an important mediator of autophagosome formation^23,24^. We found that the levels of the LC3BII/I and Beclin-1 were obviously elevated in the aorta of AD patients than in the control group by IHC and western blot (Figure 2A-B). Assessment of autophagic activity relies not only on measurements of autophagy initiation but also on degradation of autophagic substrates^25^. Therefore, we investigated the level of SQSTM1, which was apparently enhanced at the translational and transcriptional levels in AD, and this is consistent with IHC results (Figure 2C-E). In parallel, we labeled VSMCs in the damaged aorta with a VSMCs marker α-smooth muscle actin (ACTA2) to observe the localization of SQSTM1 in VSMCs. IF showed that SQSTM1 (red) co-localized with ACTA2 (green) in VSMCs from AD patients and accumulated predominantly in the middle aortic layer compared with VSMCs from normal patients (Figure 2G). We further investigated the correlation between ENO2 and LC3B or SQSTM1. Interestingly, ENO2 expression was positively correlated with LC3B expression in AD patients and similarly with SQSTM1 expression (Figure 2F). These results suggest that autophagosome clearance is impaired in AD patients, leading to autophagosome accumulation.

**Figure 2.**
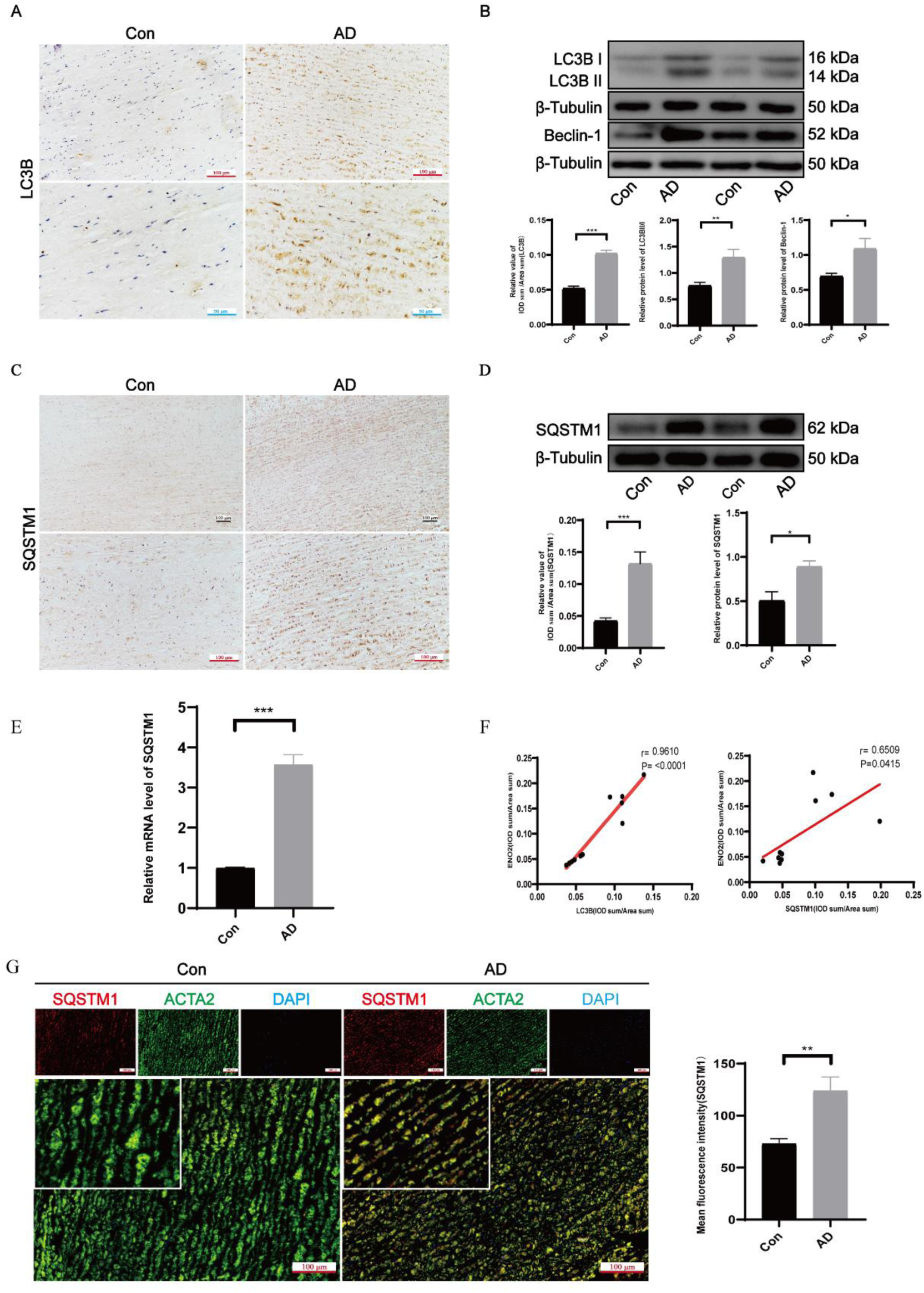
Impaired autophagosome clearance in the aorta of AD patients and a positive correlation with elevated ENO2 expression. **A** The expression of LC3B in AD patients and the control group was observed by IHC, n = 5 for each group. **B** The expression of LC3BII/I and Beclin-1 in AD patients and the control group was observed by western blot. **C-D** The expression of SQSTM1 in AD patients and the control group was observed by IHC and western blot, n = 5 for each group. **E** The mRNA expression of SQSTM1 in AD patients and the control group was observed by qRT-PCR, n=3 in each group. **F** Correlation of ENO2 with LC3B or SQSTM1 expression in human thoracic aortic tissue. **G** Representative fluorescence microscopy images of SQSTM1 (red), ACTA2 (green) and DAPI (blue) in thoracic aortic tissue from AD patients and the control group, the inset shows higher magnification images, n = 5 for each group. Data are presented as mean fold-change with error bars for SEM. T-test was used to compare the two groups of data. Statistically significant differences are indicated at * *p* < 0. 05; ** *p* < 0. 01; *** *p* < 0. 001.

### AAV9 mediates knockdown of ENO2 attenuated aortic injury and inhibits the formation of AD in mice

We next examined whether ENO2 knockdown could attenuates aortic injury and inhibits the formation of AD in mice in vivo. Pluronic gel containing AAV-GFP or AAV-ENO2 was applied to the aorta in 3-week-old male mice, and one week later, a murine AD model was established in C57B/6J mice by BAPN and Ang II (Figure 3A). Furthermore, to assess the efficiency of AAV-ENO2 infection, as expected, IF and Western blot analysis demonstrated that application of AAV-ENO2 potently inhibited ENO2 protein expression in the aorta in mice as compared with AAV-GFP at 3 weeks after transfection (Figure 3C-D). The expression of ENO2 was higher in AAV-GFP-treated AD mice aortas than in GFP-treated mice aortas (Figure S1A). Interestingly, examination of the aortas demonstrated that 76.9% of mice (10 of 13) provoked apparent mice death and contributed to the formation of AD when AD mice treated with AAV-GFP, whereas AAV-ENO2 treatment AD mice exhibited strong inhibition of the incidence of the formation of AD (4 of 13) (Figure 3B). Distinct exfoliation or rupture, degradation of elastin cleavage and excessive collagen deposition were also markedly attenuated in AAV-ENO2-treated AD mice compared with AAV-GFP-treated AD mice (Figure 3E-F). Taken together, these findings indicate that correction of ENO2 expression in AD mice by AAV-based gene therapy substantially alleviated Ang II and BAPN induced AD formation and preserved aortic structure and function.

**Figure 3.**
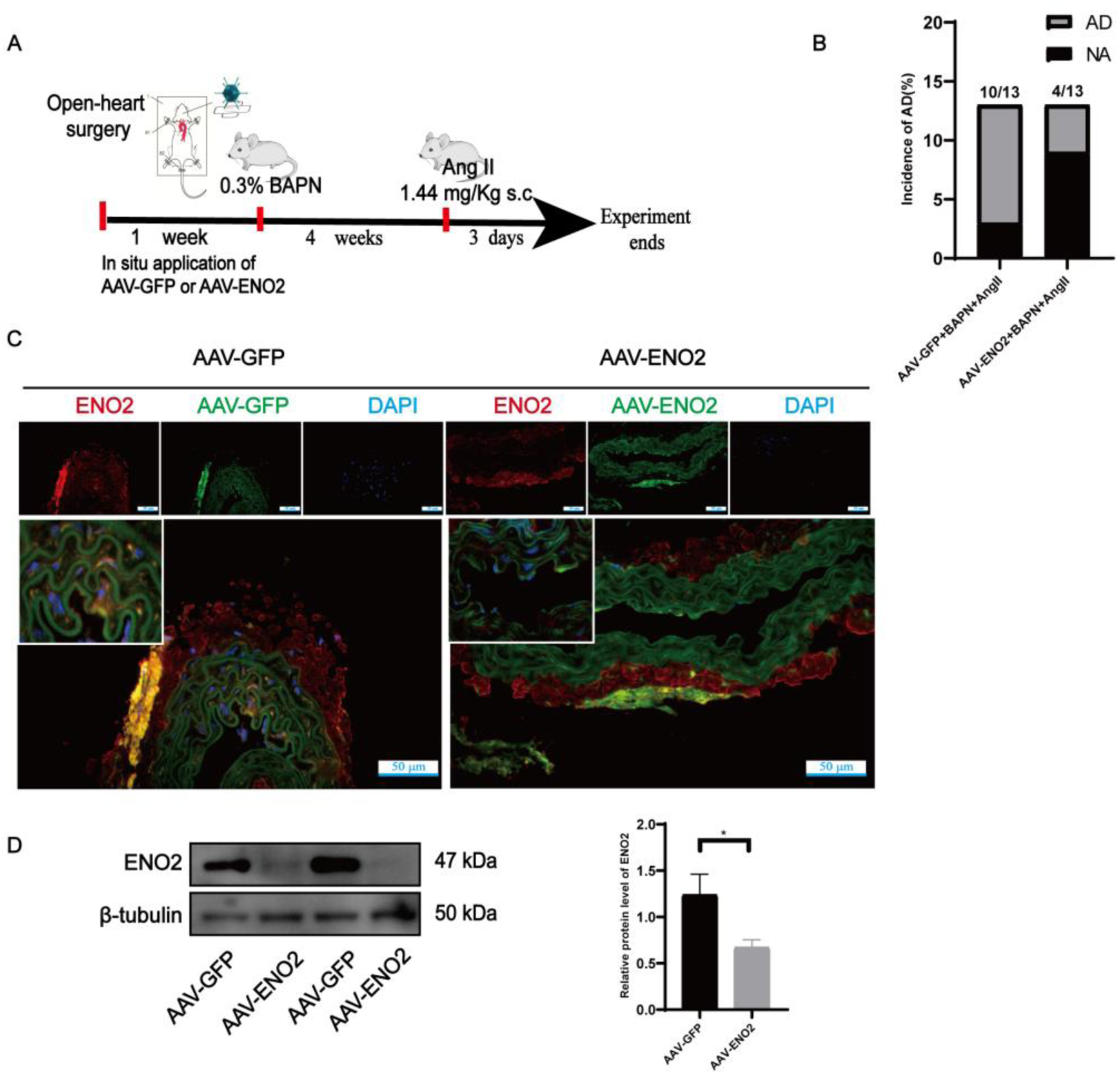
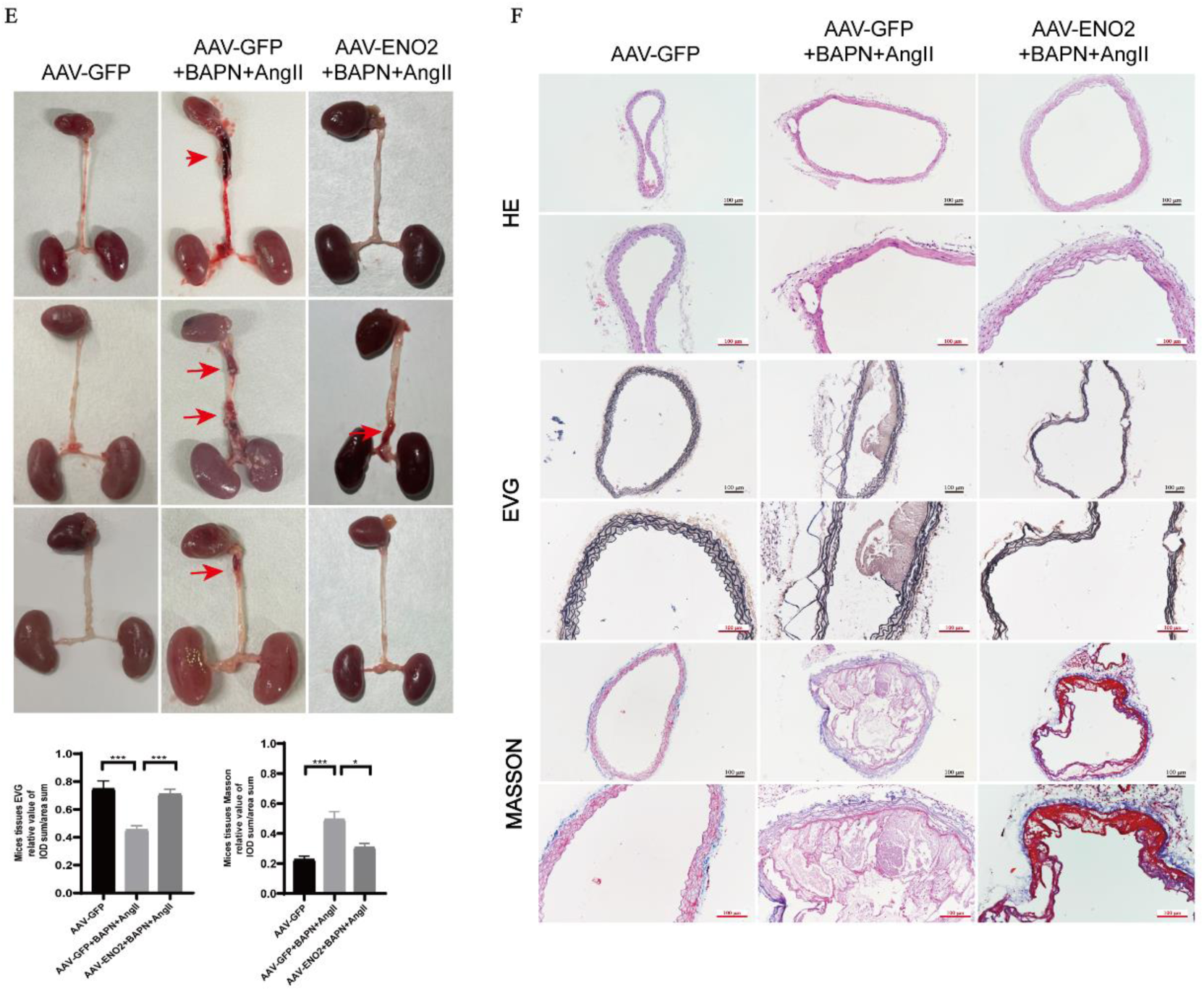
AAV9 mediates knockdown of ENO2 attenuated aortic injury and inhibits the formation of AD in mice. **A** Experimental flow chart. **B** Incidence of aortic dissection in AD mice treated with AAV-GFP or AAV-ENO2. **C-D** The expression of ENO2 by western blot and IF in the aorta of mice treated with AAV-GFP or AAV-ENO2. The inset shows higher magnification images. **E** Representative images of mouse aortic anatomy. **F** HE, EVG and Masson staining of the aortic wall of mice in each group, n≥5 in each group. Data are presented as mean fold-change with error bars for SEM. T-test was used to compare the two groups of data. One-way analysis of variance was used for data comparison among multiple groups. Statistically significant differences are indicated at * *p* < 0. 05; ** *p* < 0. 01; *** *p* < 0. 001.

### ENO2 was involved in autophagosome accumulation during AD mice and HAVSMCs stimulated with PDGF-BB

Glycolysis-related enzymes play a substantial role in autophagy, and a recent study showed that ENO2 regulates autophagy-related pathways, such as PI3K and MAPK/ERK^26^, prompting us to describe the role of ENO2 in autophagy in aortic dissection. Notably, the expression of LC3BII/I was lower in AAV-ENO2-treated AD mice aortas than in AAV-GFP-treated AD mice aortas (Figure 4A).

**Figure 4.**
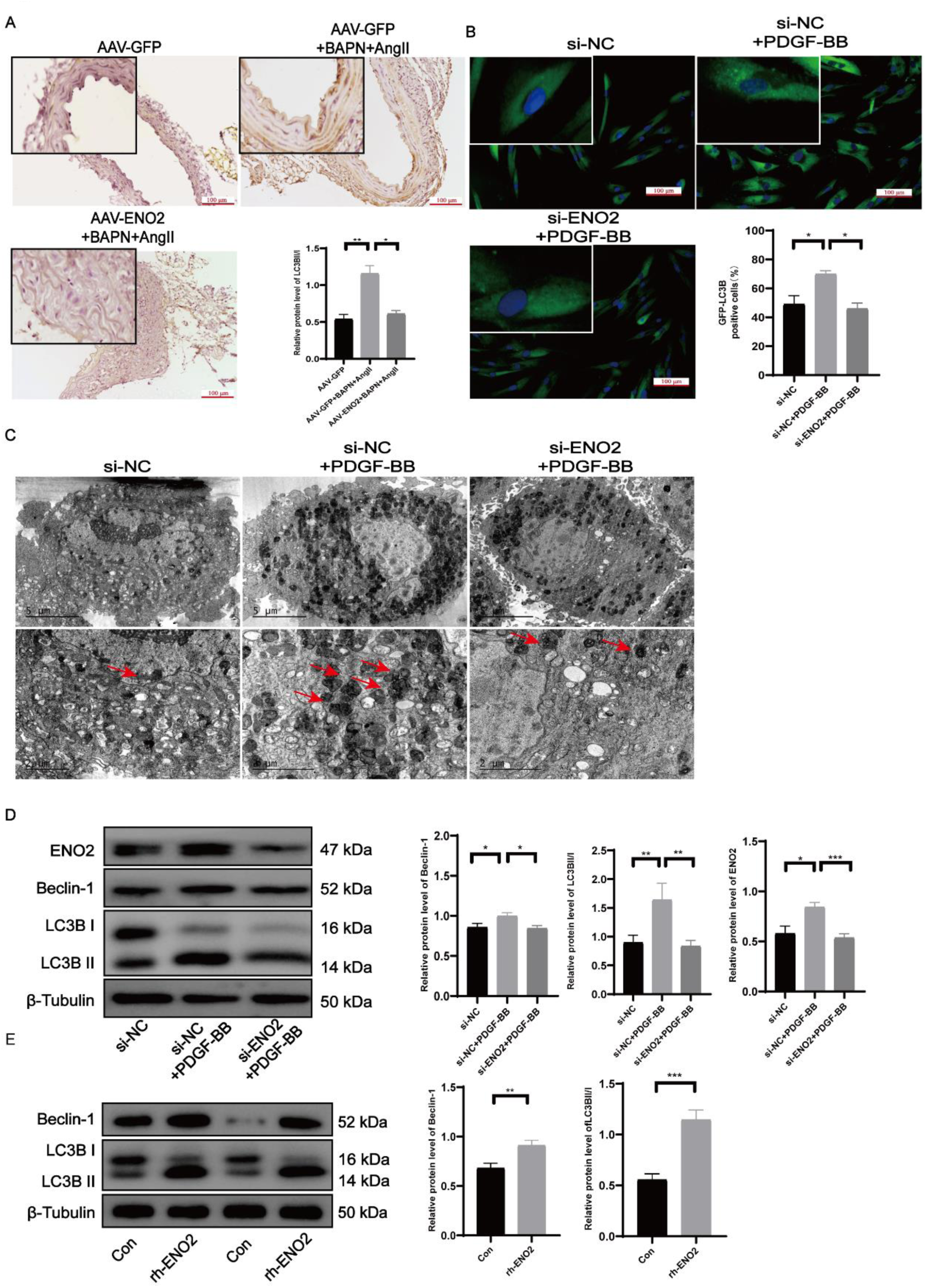
ENO2 was involved in autophagosome accumulation during AD mice and HAVSMC stimulated with PDGF-BB. **A** The expressions of LC3B was observed by IHC in the aortic wall of mice, n ≥ 5 in each group. **B** Infect HAVSMCs with Ad-GFP-LC3, followed by transfection with the indicated siRNA or treatment with the indicated chemicals. The punctate structure of GFP-LC3 was assessed by fluorescence microscopy. **C** Effect of autophagic vesicles by knockdown of ENO2 in the AD cell model. Red arrows indicate autophagic vesicles. **D** Effects on Beclin-1 and LC3BII/I expression in knockdown of ENO2 in the AD cell model. **E** Effects of increasing ENO2 level on Beclin-1 and LC3BII/I expression in HAVSMCs. HAVSMCs were treated with PDGF-BB to simulate the AD cell model. For in vitro experiments, replicate three times independently. Data are presented as mean fold-change with error bars for SEM. One-way analysis of variance was used for data comparison among multiple groups. Statistically significant differences are indicated at * *p* < 0. 05; ** *p* < 0. 01; *** *p* < 0. 001.

For confirmation, we knocked down the expression of ENO2 with three siRNAs and verified their knockdown efficiency by Western blot, and selected the most efficient si-ENO2-3 for subsequent experiments (Figure S1B). We assessed the effect of ENO2 on autophagosome accumulation. Western blot was conducted and showed that ENO2, LC3BII and Beclin-1 were significantly upregulated in HAVSMCs treated with PDGF-BB, while this effect was lost after ENO2 was knocked down (Figure 4D). In line with the Western blot results, GFP-LC3 fluorescence showed that ENO2 knockdown inhibited PDGF-BB-induced autophagosome accumulation in HAVSMCs (Figure 4B). To further investigate the effect of ENO2 on autophagosome accumulation, we observed an increase in autophagic vesicles in PDGF-BB-treated HAVSMCs by transmission electron microscopy, in contrast to the decrease in autophagic vesicles in HAVSMCs in which ENO2 was knocked down (Figure 4C). We used recombinant human ENO2 (rh-ENO2) to elevate ENO2 levels in HAVSMCs (Figure S1C). Western blot analyses indicated that the treatment of HAVSMCs with 1 μg/mL rh-ENO2 for 24 h resulted in a significant increase in Beclin-1 and LC3BII/I (Figure 4E). The above results suggest that ENO2 is involved in autophagosome accumulation in AD mice and HAVSMC stimulated with PDGF-BB.

### ENO2 impaired the autophagy flux during AD mice and HAVSMCs stimulated with PDGF-BB

We observed increased Beclin-1 and LC3BII/I expression previously detected in the aorta of AD patients, which seems to contradict previous findings of elevated SQSTM1, suggesting that the occurrence of aortic dissection may be related to disruption of autophagy flux. More importantly, we also found that knockdown of ENO2 significantly reduced autophagy accumulation in AD mice and HAVSMCs stimulated by PDGF-BB. Therefore, we speculate that ENO2 may impair autophagic flux leading to autophagosome accumulation.

To further elucidate the relationship between ENO2 and autophagic flux, we studied the expression of SQSTM1 in the aortic wall of mice by IHC. comparison to AAV-GFP-treated AD mice aortas, the expression of SQSTM1 was lower in AAV-ENO2-treated AD mice aortas (Figure 5A). Likewise, ENO2 knockdown effectively inhibited the accumulation of SQSTM1 puncta caused by PDGF-BB treatment, and reduced colocalization of these puncta with LC3B puncta (Figure 5B). These results suggest that ENO2 can reduce autophagosome accumulation and partially restore autophagy flux. Moreover, LAMP2 is a lysosome membrane protein that expresses lysosome number and together with SQSTM1 reflects lysosomal level^27^. As determined by IHC, the aorta of patients with aortic dissection exhibited decreased expression of LAMP2 compared to control subjects (Fig. S1d). Notably, we found that knockdown of ENO2 largely attenuated the inhibitory effect of PDGF-BB treatment on LAMP2 expression in HAVSMCs, thereby promoting SQSTM1 turnover (Figure 5C). In contrast, elevated ENO2 levels in HAVSMCs suppressed LAMP2 expression, resulting in SQSTM1 accumulation (Figure 5D). Altogether, these findings suggest that ENO2 may promote lysosome dysfunction and autophagosome accumulation in HAVSMCs.

**Figure 5.**
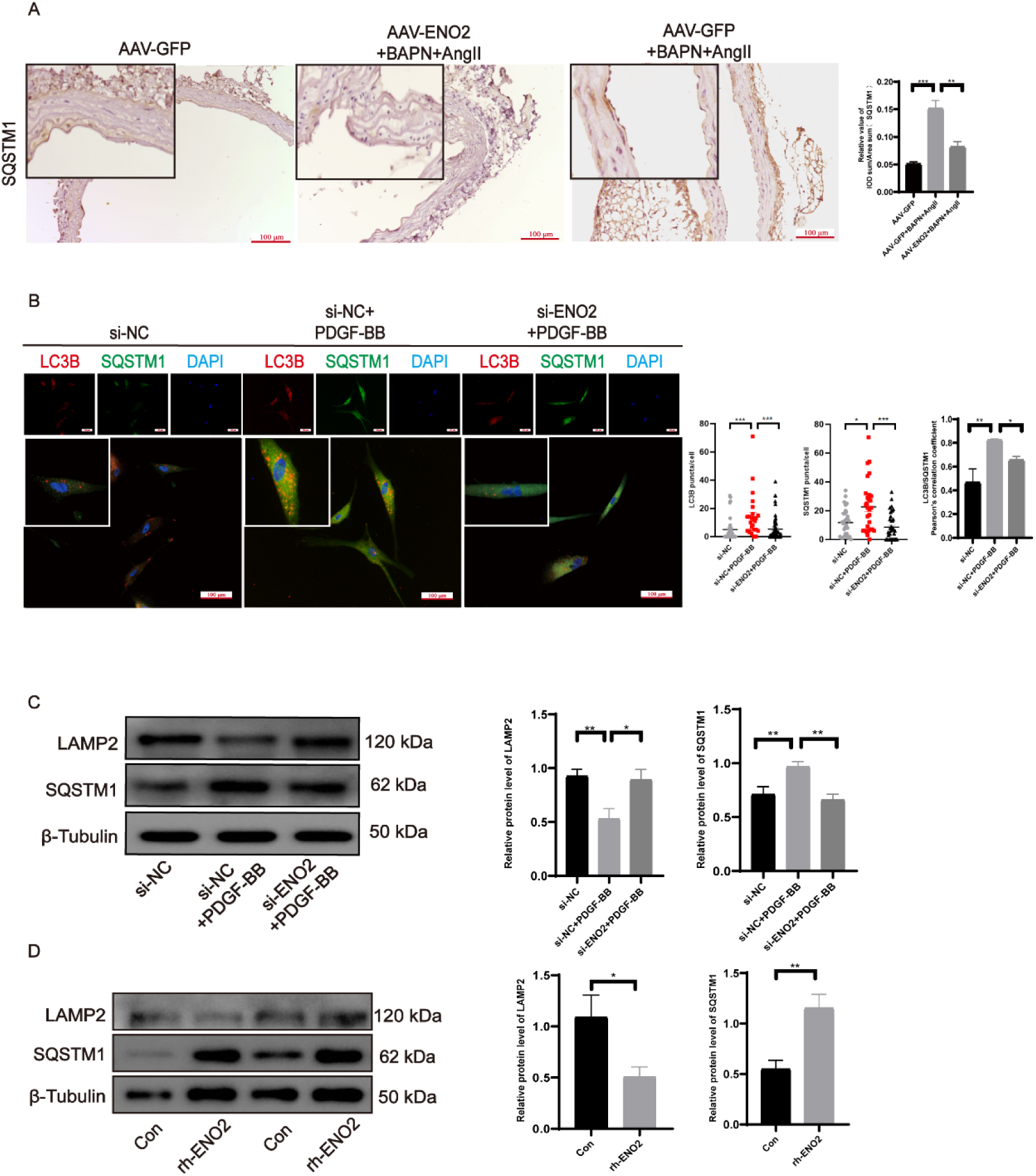
ENO2 impaired the autophagy flux during AD mice and HAVSMCs stimulated with PDGF-BB. **A** The expressions of SQSTM1 was observed by IHC in the aortic wall of mice, n≥ 5 in each group. **B** Effects on LC3B and SQSTM1 co-localization in knockdown of ENO2 by IF and Pearson’s correlation coefficients in HAVSMCs. **C-D** The effects of reduced and increased ENO2 levels on LAMP2 and SQSTM1 expression by western blot in HAVSMCs. Experiments were independently repeated three times. Data are presented as mean fold-change with error bars for SEM. One-way analysis of variance was used for data comparison among multiple groups. Statistically significant differences are indicated at * *p* < 0. 05; ** *p* < 0. 01; *** *p* < 0. 001.

### GAP43/ATF4 signaling pathway mediates the effect of ENO2 on autophagy in HAVSMCs

Next, we sought to identify the potential mechanisms by which ENO2 regulates autophagy. Growth-associated protein 43 (GAP43) is a protein, which is involved in cell proliferation and migration as well as induces apoptosis ^28^. However, its role in AD remains to be determined. There is an obvious correlation between the expression of ENO2 and GAP43. Down-regulation of ENO2 in cells leads to significantly enhanced expression of GAP43^29^, which may regulate the expression of autophagy proteins through activating transcription factor 4 (ATF4)^30–35^. Therefore, we hypothesized that ENO2 may regulate the expression of autophagy-related proteins through GAP43/ATF4 signaling pathway.

IHC and Western blot results showed that GAP43 expression was declined in the aortic tissues of AD patients (Figure 6A-B). As anticipated, the expression of GAP43 was higher in AAV-ENO2-treated AD mice aortas, comparison to AAV-GFP-treated AD mice aortas (Figure 6C). Next, we investigated whether ENO2 affected the nuclear translocation of ATF4. we performed IF to examine the effect of ENO2 on ATF4 expression and localization in HAVSMCs. Here, ATF4 staining in the nucleus was decreased in HAVSMCs with stable ENO2 down-regulation (Figure S2A). Furthermore, PDGF-BB stimulation greatly reduced GAP43 protein levels and activated ATF4 in HAVSMCs (Figure 6D). Similar results were noted for HAVSMCs treated with rhENO2 (Figure 6E). Importantly, siRNA-mediated ENO2 knockdown in PDGF-BB-treated HAVSMCs resulted in elevated GAP43 expression and inhibition of ATF4 (Figure 6D). We further found that overexpression of GAP43 with pcDNA (3. 1) in HAVSMCs treated with rhENO2 upregulated GAP43 protein and Inhibited ATF4 expression, thus reduced the expression of autophagy-related proteins Beclin-1, LC3BII/I and SQSTM1 and could restore the expression of LAMP2 (Figure 6F). We knocked down the expression of ATF4 with three siRNAs and verified their knockdown efficiency by Western blot (Figure S2B). We selected the most efficient si-ATF4-3 for subsequent experiments. Knockdown of ATF4 significantly reduced the expression of Beclin-1, SQSTM1, LC3BII/I, and upregulation the level of LAMP2 in HAVSMCs treated with rhENO2 (Figure 6G). In total, these results underline that ENO2 may impair the autophagy flux through the ENO2/GAP43/ATF4 pathway.

**Figure 6.**
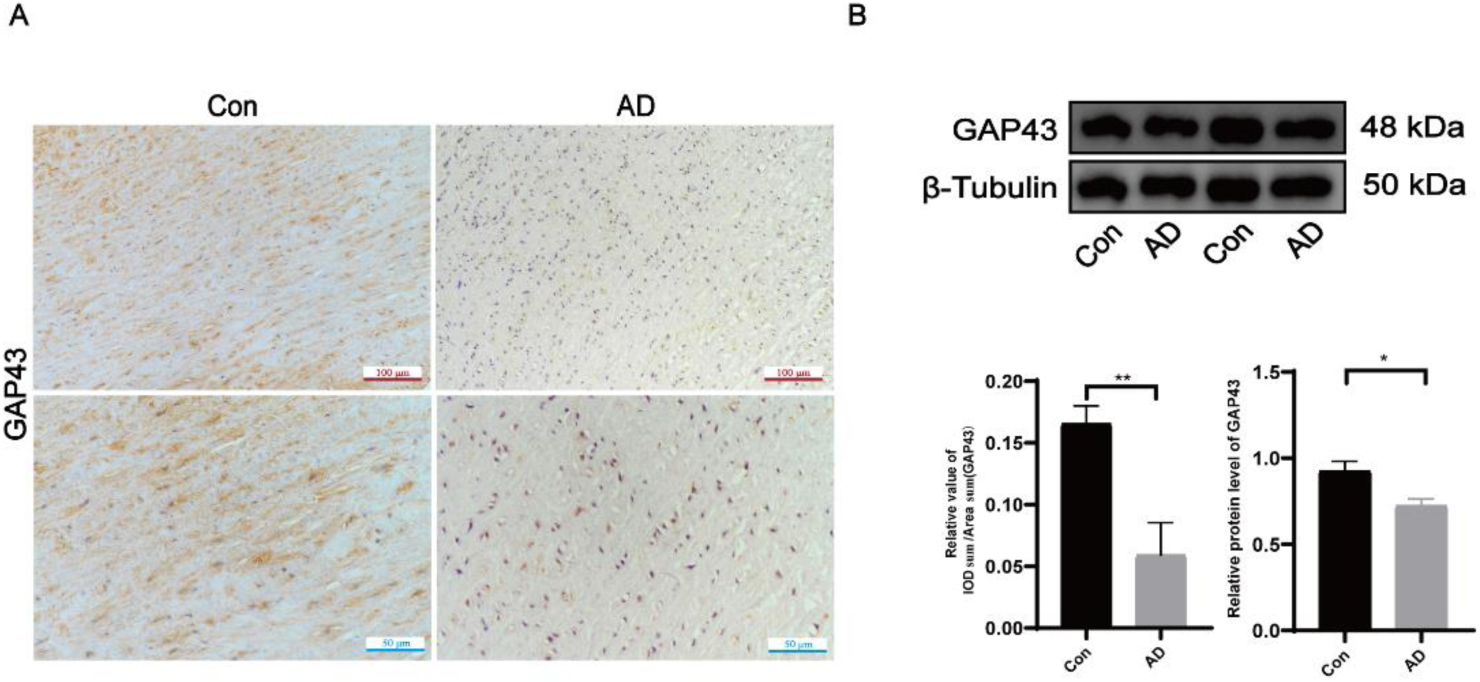
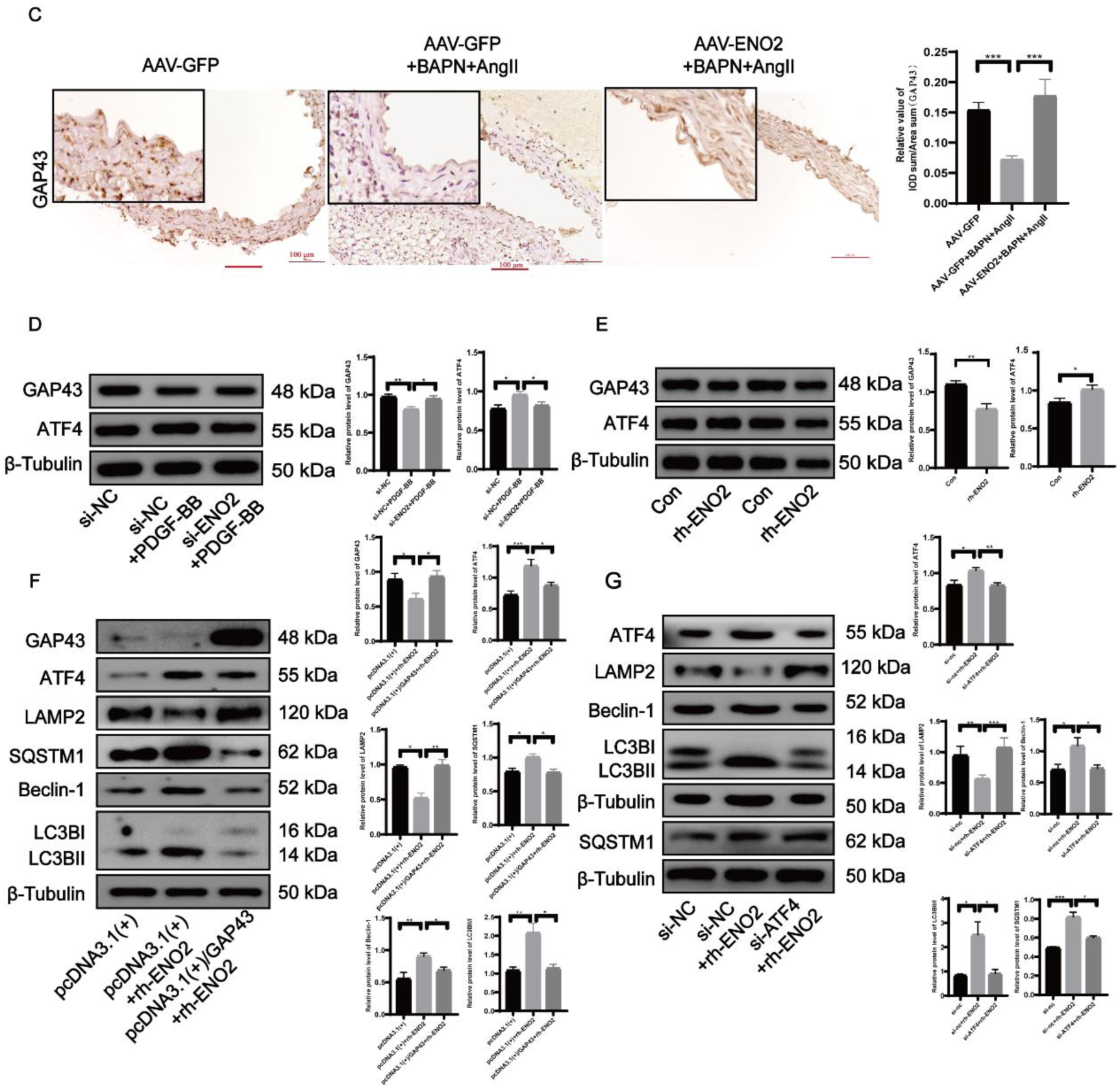
GAP43/ATF4 signaling pathway mediates the effect of ENO2 on autophagy in HAVSMCs. **A-B** The expression of GAP43 in AD patients and the control group was observed by western blot and IHC, n = 5 for each group. **C** The expression of GAP43 was observed by IHC in the aortic wall of mice, n ≥ 5 in each group. **D** In AD model cells, effect of knockdown of ENO2 on GAP43 and ATF4 expression. **E** Effect of increasing ENO2 level on GAP43 and ATF4 expression in HAVSMCs. **F** Effect of overexpression of GAP43 treated with rh-ENO2 on the expression of GAP43, ATF4, Beclin-1, LC3BII/I, SQSTM1 and LAMP2 in HAVSMCs. **G** Effect of knockdown of ATF4 treated with recombinant human ENO2 on the expression of ATF4, Beclin-1, LC3BII/I, SQSTM1 and LAMP2 in HAVSMCs. Data are presented as mean fold-change with error bars for SEM. T-test was used to compare the two groups of data. One-way analysis of variance was used for data comparison among multiple groups. Statistically significant differences are indicated at * *p* < 0. 05; ** *p* < 0. 01; *** *p* < 0. 001.

## Discussion

In this study, we demonstrated that the ENO2 protein level was upregulated in AD patients, AD mice, and PDGF-BB stimulated HAVSMCs. ENO2 upregulation during AD was related to the increased impairment of autophagic flux via the modulation of autophagosome processing and lysosome dysfunction. Likewise, ENO2 knockdown greatly reduced the development of AD, improving AD mouse survival and reducing the rupture rate of dissected blood vessels. Autophagic dysfunction with increased Beclin-1, LC3BII/I and SQSTM1 expression and decreased LAMP2 expression was noted in HAVSMCs stimulated with PDGF-BB or rh-ENO2, while ENO2 knockdown partially restored autophagic flux. ENO2 can aggravate autophagy dysfunction in HAVSMCs by regulating the expression of autophagy and lysosome proteins through a GAP43-mediated increase in ATF4 nuclear expression. These findings revealed the potential function of ENO2 during AD and provided new insights into its autophagy regulatory mechanisms.

A novel observation of this study was the critical role of ENO2 in AD via restoration of the impaired autophagic flux by facilitating autophagosome processing during AD. This role was supported by the following observations: AD patients have impaired autophagosome clearance, leading to autophagosome accumulation; ENO2 expression was positively correlated with the expression of LC3B and SQSTM1 in AD patients; autophagic dysfunction was occurred in AAV-GFP-treated AD mouse aortas; AAV-ENO2 treatment may markedly restore the blocked autophagic flux; autophagy flux was inhibited in HAVSMC stimulated with PDGF-BB or rh-ENO2, which showed increased expressions of Beclin-1, LC3BII/I, and SQSTM1, decreased expression of LAMP2, while autophagy flux was partially restored by ENO2 knockdown. Consistently, ENO2 was critical in the restoration of impaired Autophagic Flux during AD mice and HAVSMCs stimulated with PDGF-BB.

VSMCs are a principal cell type of the aorta and play a crucial role in the pathogenesis of AD in humans^36^. Upon vascular injury, such as atherosclerosis, aortic dissection, and aneurysm, VSMCs dedifferentiate from a “contractile” phenotype to a highly “synthetic” phenotype characterized by an enhanced rate of glycolytic flux ^37–41^. However, enhanced glycolysis may lead to the inhibition of autophagosome degradation ^42^. As a key enzyme in glycolysis, ENO2 is directly involved in glycolysis, catalyzing the reversible conversion of 2-phosphoglycerate to phosphoenolpyruvate^43^. Dysregulation of ENO2 expression is associated with lung injury, endothelial cell migration, lymphocyte apoptosis, and the MAPK signaling pathway ^44^. In this study, we found that ENO2 expression levels were appreciably elevated in AD patients. The increase in glycolytic flux in the aorta of AD tissue, which leads to the upregulation of the key glycolytic enzyme ENO2, may be an injury mechanism of VSMCs during the progression of AD, and this injury mechanism might partly lead to the dysfunction of autophagy in AD.

Accumulating evidence indicates that GAP43 seems to be beneficial in cardiovascular disease^45^. It is a PKC-activated protein that is crucial for apoptosis as well as epithelial mesenchymal transition^28^. GAP43 may regulate autophagy gene translation through the transcription factor ATF4^30–35^. Our data showed that GAP43 expression levels were lower in the aortic tissues of AD patients than in normal tissues. More importantly, the aortic media of AAV-ENO2-treated AD mice upregulated GAP43 protein expression and enhanced ATF4 activation was accompanied by impaired autophagic flux compared with the aortic media of AAV-GFP-treated AD mice. Consistently, in HAVSMCs treated with PDGF-BB, knockdown of ENO2 resulted in a significant increase in GAP43 expression, inhibition of ATF4 expression, and suppression of ATF4 translocation to the nucleus. Similarly, increased levels of ENO2 in HAVSMCs were able to reduce GAP43 level and promote ATF4 expression. Thus, the GAP43/ATF4 pathway is involved in the aortic effects of ENO2 by regulating autophagic flux in AD mice and HAVSMCs stimulated with PDGF-BB.

Furthermore, overexpression of GAP43 abolished exogenous ENO2-mediated impairment of autophagosome clearance and activation of ATF4. ATF4 is essential for the transcriptional activation of genes responsible for autophagosome formation, elongation and function, and is involved in the induction of autophagy genes such as Beclin-1, LC3B and SQSTM1 and the downregulation of LAMP2 to regulate autophagy and lysosome-related protein synthesis to balance cellular homeostasis and improve survival stress ^31–35^. Notably, knockdown of ATF4 rescued exogenous ENO2-increased autophagic flux impairment in HAVSMCs, therefore ATF4 acts as a negative regulator of autophagic flux. In addition, ATF4 can induce CHOP expression thereby responding to ER stress, leading to SMC apoptosis and promoting TAAD formation^46^. It seems to play multiple roles in AD. Together, these findings suggest that the ENO2-regulated GAP43/ATF4 pathway plays an essential role in the regulation of autophagic flux in AD mice and in HAVSMCs stimulated with PDGF-BB.

Another finding was the important role of ENO2 in impeding the development of AD. This finding was supported by the following observations: ENO2 was expressed in the aortic media^47^ and was abnormally elevated in the aorta of AD patients. Knockdown of ENO2 attenuated aortic lesions in AAV-ENO2-treated AD mice compared with AAV-GFP-treated mice. Knockdown of ENO2 reduced the incidence of AD and decreased mortality in AAV-ENO2-treated AD mice compared with AAV-GFP-treated mice. In addition, we found that a gel containing adenovirus was applied to the aorta during open-heart surgery in mice to achieve better infection and to conserve adenovirus dosage.

Numerous studies have shown that inhibition of autophagic flux can lead to vascular injury, for example by inhibiting the proliferative migration and contractile phenotype of HAVSMCs, as well as increasing vascular senescence and calcification^48–50^. Likewise, defective autophagy in VSMCs enhances the incidence and severity of AD. These data suggest that autophagy of VSMCs is essential for maintaining vascular integrity during the development and progression of AD. This interconnection between ENO2 and the autophagy pathway is important in causing vascular injury and deserves further consideration.

Since ENO2 might be expressed in other cell types of mouse’s aorta, the effects of ENO2 in vivo are probably not exclusively derived from HAVSMCs. Thus, our cellular experimental manipulation results suggest, but do not confirm, an in vivo role of ENO2 in impaired autophagic flux in the aorta. The possibility that other cells expressing ENO2 are involved in regulating impaired aortic autophagic flux warrants further investigation. In addition, the clinical sample size is small, so more work is needed on how to define thresholds and how to transfer them to clinical use.

Other key glycolytic enzymes have been documented to play an important role in AD^51,52^. We are currently preparing to build an early warning system for AD by combining well-defined clinical risk factors, changes in the levels of key enzymes of glucose and lipid metabolism and related factors. Subsequently, we validate the reliability and sensitivity of early screening warning systems in cardiovascular disease populations, and animal models for calibration and application. The AD early screening and early warning system can be applied to cohorts with a population size of 200,000, screening out high-risk AD groups for early intervention.

Taken together, we report here that upregulation of ENO2 exacerbates autophagic dysfunction through the GAP43/ATF4 pathway. These findings also suggested the importance of maintaining autophagic homeostasis to limit the development of AD.

## Nonstandard Abbreviations and Acronyms

AD, Aortic dissection; ENO2, Enolase 2; GAP43, Growth-associated protein 43; ATF4, Activating transcription factor 4; SQSTM1, Sequestosome 1; LAMP2, Lysosome membrane protein 2; LC3B, Microtubule-associated protein light chain 3B; HAVSMCs, Human aortic vascular smooth muscle cells; Rh-ENO2, Recombinant human ENO2 protein, Adeno-Associated Virus (AAV)

## Author contributions

**Liangwan Chen and Yumei Li:** Designing the study. **Weixing Cai and Li Zhang:** Writing - original draft. **Jing-Jing Lin, Qiuying Zou and Chaoyun Wang:** Conducting experiments. **Xi Yang, Xiaohui Wu, Jianqiang Ye and Hui Zheng:** Data analysis. **Lin Zhang, Lin Zhong, Xinyao Li, Keyuan Chen and Jiangbin Wu:** Reviewing and editing the manuscript.

## Study approval

The study was conducted in accordance with the ethical guidelines of the Declaration of Helsinki and approved and conducted under the guidance of the Ethics Committee of Fujian Medical University Union Hospital (No. 2019-36). All subjects signed a written informed consent form. The experimental animal ethics committee of Fujian Medical University (IACUC FJMU-Y-0763) approved that the experimental design and implementation of this study were in accordance with animal welfare and ethical requirements, and passed the ethical review of animal experiments.

## Acknowledgements

The authors thank the Union Hospital of Fujian Medical University and the Experimental Animal Center of Fujian Medical University for their support of the experiments. The authors thank the patients and medical staff for their support of this

